# A Bayesian Regularized and Anotation-Informed Integrative Analysis of Cognition (BRAINIAC)

**DOI:** 10.1101/2023.07.24.550424

**Authors:** Rong Zablocki, Bohan Xu, Chun-Chieh Fan, Wesley K. Thompson

## Abstract

Here we present a development of the novel Bayesian Regularized and Anotation-Informed Integrative Analysis of Cognition (BRAINIAC) model. BRAINIAC allows for both estimation of total variance explained by all features for a given cognitive phenotype, as well as a principled assessment of the impact of annotations on relative enrichment of features compared to others in terms of variance explained, without relying on a potentially unrealistic assumption of sparsity of brain-cognition associations. We apply the BRAINIAC model to resting state fMRI data from the Adolescent Brain Cognitive Development Study.

## 2. Introduction

Functional Magnetic Resonance Imaging (fMRI) measures changes in blood oxygenation and blood flow related to neuronal activity, providing researchers with the means to study human brain function *in vivo* [8]. fMRI is safe, relatively inexpensive, and has fairly good spatial and temporal resolution. Over the past two decades fMRI has become an essential and widely-used tool for assessing the neural substrate of human behavior and psychiatric disorders.

However, in recent years neuroimaging research has experienced a “replication crisis” [2], wherein many initial published positive findings do not replicate in subsequent research. It has been hypothesized that this crisis has been caused by an unfortunate combination of factors, including small effect sizes, inadequate sample sizes, high-dimensional feature spaces (with a correspondingly high researcher degree of freedom) and publication bias towards ”significant” associations. In support of this, it has been recently shown that, if estimated from studies with large samples, effects sizes of brain-behavior relationships are often considerably smaller than has been published in the past [3, 5]

Furthermore, it appears that brain-behavior associations, rather than being localized to a sparse set of regions, are widely distributed across the brain. For example, Zhao et al. [14] found that whole-brain ridge regression of all vertices outperformed methods (e.g., LASSO) that assumed sparsity for predicting cognitive ability using the N-back task fMRI data in the Adolescent Brain Cognitive Development (ABCD) Study. This scenario argues for using statistical methods that integrate unthresholded effects across the whole brain (e.g., all vertices, voxels, or networks). This is precisely the scenario that Genome-Wide Association Studies (GWAS) encountered in the last decade [6]. Candidate gene studies failed to replicate, individual effect sizes were smaller than researchers had previously believed for gene-behavior associations, and widely spread across the genome.

In response to this, Yang et al. [13] proposed a variance-components analysis algorithm, termed Genome-wide Complex Trait Analysis (GCTA) for GWAS data. GCTA is designed to assess the total fraction of variance explained (FVE) for all loci in a GWAS, and can be applied when the number of loci far exceeds the number of subjects. GCTA requires subject-level data to fit. Other FVE methods, including LD-Score Regression [1] and GWASH [11], work on the GWAS *summary statistics* typically consisting of publicly-released regression coefficients and p-values for each genetic locus.

The statistical model employed by GCTA is as follows. Let ***y*** = (*y*_1_, …, *y*_*n*_)^*T*^ be a mean-zero vector of standardized continuous phenotype values on *n* independent participants. Let **X** denote an *n* × *B* predictor matrix

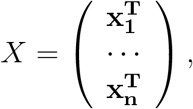

where the **x**_**i**_ are *B*-dimensional vectors of features (here, Single Nucleotide Polymorphisms, or SNPs). The columns of ***X*** are standardized to have zero mean and unit variance. We assume the model

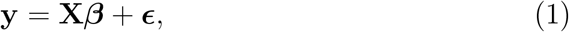

where ***β****∼* N(**0, Σ**_*β*_) independently from 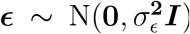. GCTA further assumes that 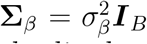, where ***I***_*B*_ is the *B × B* identity matrix. If *y* has also been standardized so that Var(*y*) = 1, the FVE by ***X*** for phenotype ***y*** is given by

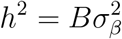

The paramteter *h*^2^ is called the *chip heritability* in the GWAS literature (the fraction of the phenotypic variance explained by additive linear effects of a specific set of genetic loci). The GCTA model is fitted using Restricted Maximum Likelihood (REML) implemented with the average information (AI) algorithm [13].

Once the model parameters are estimated, the random effects can be obtained via the Best Linear Unbiased Predictor (BLUP) for ***β***. The BLUP is given by

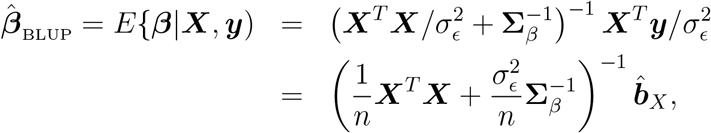

where 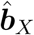 = ***X***^*T*^ ***y****/n* is the *B*-dimensional vector of mass univariate linear regression coefficients of **y** on **X**. For a new subject’s genetic data, denoted by ***x***new, the predicted phenotype from the BLUP is

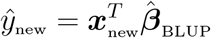

The GCTA model (1) assumes all ***β*** are independent and exchangeable, i.e 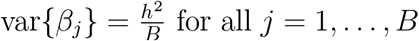 for all *j* = 1, …, *B*. However, this assumption ignores any information that researchers may have about the relative importance of some features compared to others. As a consequence, the standard GCTA model is minimally informative about which features are more or less important for predicting ***y***, and hence not fundamentally geared to test the relative importance of features in predicting the phenotype of interest.

In this paper we describe a Bayesian hierarchical variance components model developed for application to fMRI data, in which the number of features can be much larger than the number of individuals. The goals of the model are 1) to estimate the FVE of all imaging features; and 2) to assess the relative enrichment of variance explained for individual features based on feature-specific “annotations”. Annotations can be multi-dimensional, and of mixed type (i.e., discrete and/or continuous). It is hoped that by expanding the variance components model (1) to allow for feature annotations, that the model will enable expert input and hence become more useful for researchers to test hypotheses about which factors are more or less related to explaining brain-behavior associations.

## 3. Methods

### 3.1 BRAINIAC Model

Here we present a development of the novel Bayesian Regularized and Anotation-Informed Integrative Analysis of Cognition (BRAINIAC) model. BRAINIAC allows for both estimation of total variance explained by all features for a given cognitive phenotype, as well as a principled assessment of the impact of annotations on relative enrichment of features compared to others in terms of variance explained, without relying on a potentially unrealistic assumption of sparsity of brain-cognition associations.

Let ***Y*** denote a vector of behavioral phenotypes collected from *n* participants in a brain-wide association study (BWAS). Let ***X*** be the corresponding *n× N* matrix of fMRI features. We assume that ***Y*** and the columns of ***X*** have been standardized to have mean zero and unit variances. Similar to GCTA, the proposed model is

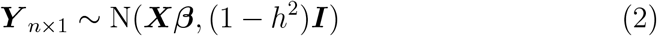

where ***β*** is a *N* -dimensional vector of regression parameters of per-feature effect sizes and 0 *< h*^2^ *<* 1 represents the fraction of the variance explained (FVE) by all features, so that 1-*h*^2^ is the variance unexplained by the model features. We propose a normal prior for **β**

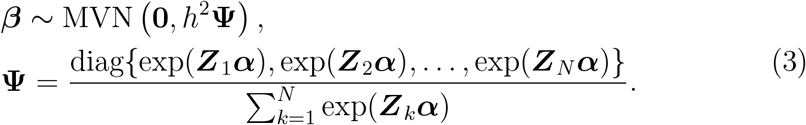

The variance of ***β*** is comprised by two components: *h*^2^ quantifies the average variance explained across all features, whereas the *N×N* diagonal matrix **Ψ** quantifies how annotations ***Z*** modulate the effect size variance of individual features. Here, ***Z*** = (***Z***_1_, ***Z***_2_, …, ***Z***_*N*_)^*T*^ is a *N×M* matrix, where *M* is the number of the annotations per feature and ***α*** is the M-vector parameter. The martrix **Ψ** is scaled to ensure the average variance of ***β*** is *h*^2^ across all features. If ***α*** = **0**, the prior of ***β*** reduces to MVN 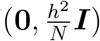. On the other hand, when **α***≠* **0**, the variance is heteroscedastic and some features account for more variance than others depending on the levels of ***Z***_***k***_.

To complete the Bayesian model, we assign a diffuse normal prior to ***α***

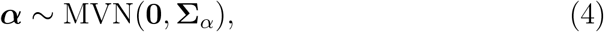

whereas *h*^2^ is given a Beta prior defined on the interval [0,1],

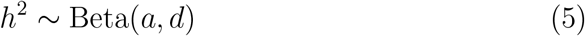

where *a* and *d* are hyperparameters; if they are both set to be 1, the Beta distribution becomes a Uniform distribution on interval [0,1]. Bayesian inference is performed via a Markov Chain Monte Carlo (MCMC) sampling algorithm, implemented in Python and described in Supplementary Section (9.1) and (9.2). The Python code is publicly available.

## 4. Results

### 4.1 Application to Resting-State fMRI Data

The BRAINIAC model was applied to data taken from the Adolescent Brain Cognitive Development (ABCD) Study, using resting-state fMRI data from Release 4.0. The ABCD Study has primary data collections sites at 21 research institutes across the USA. It is a longitudinal study of brain development and child health in the United States; *n* = 11,880 children aged 9-10 and their parents/guardians were recruited into the study at baseline, and their biological and behavioral development will be measured and followed up through adolescence into young adulthood. Details of the study design are given in **Garavan et al**., **Dick et al**. Details of the fMRI data collection and processing ate given in **Hageler et al**..

The resting-state MRI features of interest, ***X***, are “connectivities” (Fisher z-transformed Pearson correlations) between pairs of cortical vertices. The cognitive phenotype ***Y*** is the the crystalized intelligence score measured by the NIH Toolbox **Thompson et al**.. We used resting state MRI and NIH Toolbox data collected at either the baseline visit or the year 2 follow up visit. Specifically, we merged ***X*** and ***Y*** and retained participants without any missing values. If a participant only had baseline data, we included the baseline; if he or she only had 2-year visit data, included the 2-year visit; if he or she had both, we included the baseline data. In other words, we included the earliest non-missing features and phenotype, such that each remaining participant in the study had at most one row in ***X*** and a corresponding entry in ***Y***. The final sample size *n* = 9,323. The number of features (pairwise correlations) was *N* = 46,360, yielding an approximately 5:1 ratio of features *N* to participants *n*.

To regress out any potential influence from potenital confounders, wer inluded age, sex, the first 10 principal components of genetic ancestry (PC1 to PC10), and MRI scanner instance. Letting *w* denote the number of covariates, we denote by ***W*** the *n* (*× w* + 1) covariate matrix (including intercept). Residualized ***X***_*r*_ equals ***X W***− (***W***^*T*^ ***W***)^*−*1^***W***^*T*^ ***X*** and ***Y*** _*r*_ equals ***Y***−***W*** (***W***^*T*^ ***W***)^*−*1^***W***^*T*^ ***Y***. the residualized ***X***_*r*_ and ***Y***_*r*_ were then standardized to have zero column means and unit variances.

The annotations ***Z*** we used in these analyses were the assignment of cortical vertices into network labels, using the 13 resting state networks described in **Dosenbach et al**. Each pair of cortical vertices forms an ”edge” with edge strength given by the residualized Fisher Z-transformed Pearson correlation of activation from the resting state MRI data. The percentages of edges in each of these annotations annotations is displayed in Supplementary Section (9.4) Table (4) and Table (5).

**Table 1:** Estimates^1^ from single-annotation model with 13 within-network annotations (phenotype : nihtbx cryst uncorrected)

| Annotations | $\hat{\alpha}$ | $\hat{h}^2$ | $\hat{R}^2$ |
| --- | --- | --- | --- |
| Auditory-Auditory | 0.06 (-0.35,0.15) | 0.32 (0.30,0.35) | 0.30 (0.28,0.32) |
| CinguloOperc-CinguloOperc | -0.28 (-0.83,0.004) | 0.32 (0.30,0.35) | 0.30 (0.28,0.32) |
| CinguloParietal-CinguloParietal | 0.05 (0.02,0.07) | 0.32 (0.29,0.35) | 0.30 (0.28,0.32) |
| Default-Default | 0.15 (0.05,0.24) | 0.32 (0.29,0.35) | 0.30 (0.28,0.32) |
| DorsalAttn-DorsalAttn | -0.33 (-0.83,0.08) | 0.32 (0.30,0.35) | 0.30 (0.28,0.32) |
| FrontoParietal-FrontoParietal | 0.05 (-0.23,0.15) | 0.32 (0.30,0.35) | 0.30 (0.28,0.32) |
| RetrosplenialTemporal-RetrosplenialTemporal | -0.15 (-0.62,0.02) | 0.32 (0.30,0.35) | 0.30 (0.28,0.32) |
| SMhand-SMhand | 0.05 (-0.48,0.15) | 0.32 (0.29,0.35) | 0.30 (0.28,0.32) |
| SMmouth-SMmouth | -0.24 (-0.72,0.02) | 0.32 (0.30,0.35) | 0.30 (0.28,0.32) |
| Salience-Salience | -0.27 (-0.61,0.02) | 0.32 (0.30,0.35) | 0.30 (0.28,0.32) |
| Subcort-Subcort | -0.21 (-0.50,0.03) | 0.32 (0.30,0.35) | 0.30 (0.28,0.32) |
| VentralAttn-VentralAttn | 0.07 (-0.17,0.14) | 0.32 (0.29,0.35) | 0.30 (0.28,0.32) |
| Visual-Visual | -0.03 (-0.39,0.12) | 0.32 (0.30,0.35) | 0.30 (0.28,0.32) |

<sup>1</sup>Estimates ( $\hat{\alpha}$ , $\hat{h}^2$ , $\hat{R}^2$ ) shown are the overall median value and 95% posterior credible interval (CI) from the intermix of 6 chains per annotation.
$\hat{R}^2 = \text{var}(\mathbf{X}_r\hat{\beta})/\text{var}(\mathbf{Y}_r)$ is calculated for each chain at each collected iteration for all 6 chains per annotation, the overall median and 95% CI per annotation are reported. $\hat{\beta}$ is a vector estimate of $\beta$ .

**Table 2:** Estimates^1^ from multiple-annotation model with 2 within-network annotations (phenotype : nihtbx cryst uncorrected)

| Annotations | $\hat{\alpha}$ | $\hat{h}^2$ | $\hat{R}^2$ |
| --- | --- | --- | --- |
| CinguloParietal-CinguloParietal | 0.05 (0.02,0.07) | 0.32 (0.29,0.34) | 0.30 (0.28,0.32) |
| Default-Default | 0.15(-0.04,0.23) |  |  |

<sup>1</sup>Estimates ( $\hat{\alpha}$ , $\hat{h}^2$ , $\hat{R}^2$ ) shown are the overall median value and 95% posterior credible interval (CI) from the intermix of 6 chains. In multiple-annotation model, only one set of estimates for $\hat{h}^2$ and $\hat{R}^2$ , respectively.
$\hat{R}^2 = \text{var}(\mathbf{X}_r\hat{\beta})/\text{var}(\mathbf{Y}_r)$ is calculated for each chain at each collected iteration for all 6 chains, the overall median and 95% CI per annotation are reported. $\hat{\beta}$ is a vector estimate of $\beta$ .

We first applied these annotations to the BRAINIAC model one at a time; in other words, for each run *M* = 1 (single-annotation model). For each model, we ran 6 MCMC chains invoked simultaneously per annotation, each chain was run for 10,000 iterations with 3,000 burn-in period and thinning of 10.

Posterior parameter estimates of single-annotation model from between-network and within-network are presented in Table (**??**) and Table (1), respectively. Estimates shown are the overall median value and 95% posterior credible interval (CI) from the intermix of 6 chains per annotation. Although 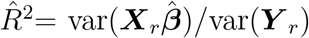 is not a parameter in the Bayesian model, it is the variance explained by the features from the regression and it is closely related to *h*^2^. The denominator of 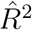 can be ignored when ***Y*** _*r*_ is standardized. 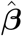 is a vector estimate of **β**.

The next step was to re-run the model using all of the annotations (***α***) from single-annotation model with posterior 95% CIs that did not include zero in their 95% posterior credible intervals and re-run the subset of the annotations in the same model (*M >* 1, multiple-annotation model). Table displays the results of multiple-annotation models from between-network and within-network, respectively.

As can be seen in these two Tables, two within-network annotations (Cingulo-Parietal and Default Mode) were significantly enriched for associations (Cingulo-Parietal ***α*** = 0.05 and Default Mode ***α*** = 0.15).

## 5. Discussion

Here, we have developed the BRAINIAC model for analyzing whole-brain data associations with cognitive phenotypes. This model allows for estimation of the overall variance explained by all imaging features simultaneously, while also allowing for feature-sepcific difference in the amount of variance explained. While we applied the BRAINIAC model to resting state MRI associations with crystalized intelligence from the ABCD Study, the model is quite general and could be applied to a number of different modalities (e.g., task-based MRI), outcomes (e.g., psychiatric symptoms), and annotations (e.g., gene expression in the human brain). In future work we plan on implementing the BRAINIAC model in a more computationally efficient algorithm (e.g., Hamiltonian MCMC or Variational Bayes) and incorporating a broad variety of annotations to furhter investigate brain-behavior relationships.

## 6 Acknowledgements

Data used in the preparation of this article were obtained from the Adolescent Brain Cognitive Development (ABCD) Study (https://abcdstudy.org), held in the NIMH Data Archive (NDA). This is a multisite, longitudinal study designed to recruit more than 10,000 children aged 9–10 and follow them over 10 years into early adulthood. The ABCD Study® is supported by the National Institutes of Health and additional federal partners under award numbers U01DA041048, U01DA050989, U01DA051016, U01DA041022, U01DA051018, U01DA051037, U01DA050987, U01DA041174, U01DA041106, U01DA041117, U01DA041028, U01DA041134, U01DA050988, U01DA051039, U01DA041156, U01DA041025, U01DA041120, U01DA051038, U01DA041148, U01DA041093, U01DA041089, U24DA041123, and U24DA041147.

A full list of supporters is available at https://abcdstudy.org/federal-partners.html. A listing of participating sites and a complete listing of the study investigators can be found at https://abcdstudy.org/ consortium members. ABCD consortium investigators designed and implemented the study and or provided data but did not necessarily participate in the analysis or writing of this report. This manuscript reflects the views of the authors and may not reflect the opinions or views of the NIH or ABCD consortium investigators.

## 7. Author Contributions

WKT formulated the BRAINIAC model. RZ derived the conditional posteriors and ran the data simulations. BX ran the real data analyses. CF provided guidance on the real data analsyes and interpretation of results. All co-authors participated in drafting the manuscript.

## 8. Tables

## 9. Supplementary Materials

### 9.1 Conditional Posteriors and Gibbs Sampling Algorithm

Let 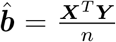 denote the marginal least squares (MLS) effect size estimates and 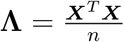 denote the in-sample correlation matrix between features.

The full conditional distributions for parameter estimates of **β**, *h*^2^ and ***α*** are described below. In the posterior inference, *n* is the number of the subjects, *N* is the number of the features, *M* is the number of annotations. The data ***X*** and ***Y*** are replaced by sufficient statistics 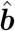 and **Λ**.

#### 9.1.1 Posterior *β*

Posterior conditional distribution of ***β*** has a closed form of MVN and can be drawn from Gibbs sampler directly:

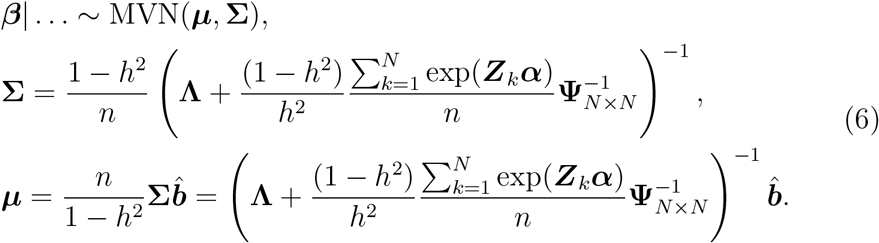

#### 9.1.2 Computational feasibility: block partition

Due to large dimension of **Λ** (*N × N*), ***β*** can be drawn block by block in a way similar to partitioned regression. The partition of the features doesn’t have to be scientifically meaningful and blocks don’t have to be independent to each other. The main idea is to split large number of features into relatively smaller blocks for computational feasibility. Let *j* = 1, 2, …, *J* (*J* is the total number of the blocks) be the index of the block, *j*^*∗*^ be the complement blocks other than *j* and *N*_*j*_ is the total number of features within *j*^*th*^ block, then the block-wise conditional distribution of **β**_*j*_ is

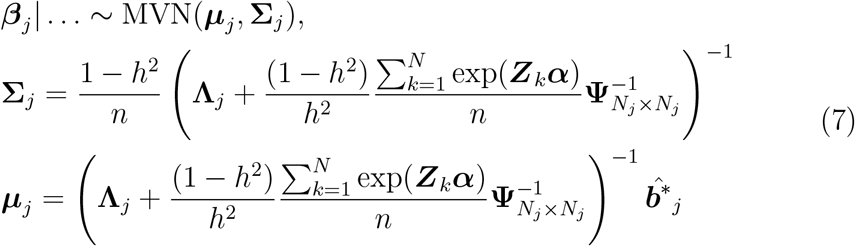

where 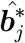 is the adjusted MLS estimate of 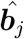derived as

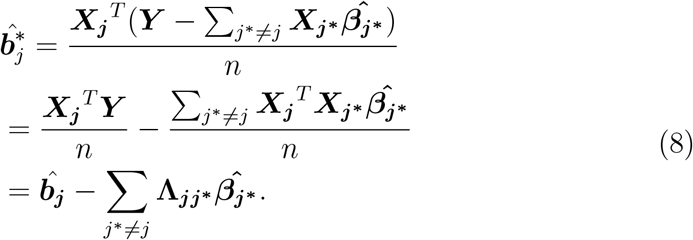

The adjustment is to remove the possible effect contributed from out-*j* blocks. The estimated effect size 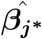 can be obtained during MCMC iterations.

#### 9.1.3 Posterior *h*^2^

Posterior *h*^2^ doesn’t have a closed form and its conditional distribution is proportional (symbolized by *∝*) to

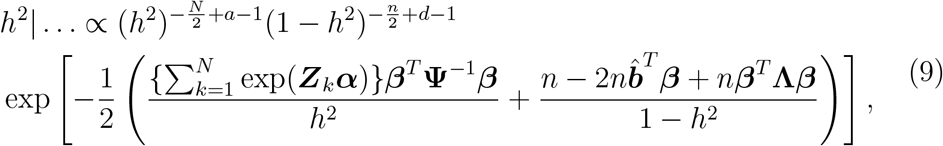

*h*^2^ is drawn via Griddy-Gibbs Sampler (GGS, [10]).

#### 9.1.4 Posterior α

Posterior ***α*** doesn’t have a closed form, either, and its full conditional distribution is proportional to

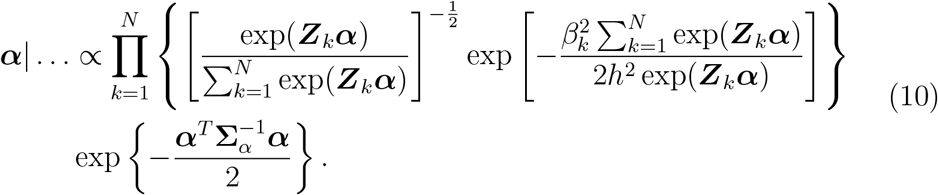

Since annotations are assumed to be independent to each other, hence, ***α*** can be drawn one by one via Metropolis-Hastings (MH) sampler ([4, 7]). Let *m* = 1, 2, …, *M*, the corresponding equation (10) with respect to (w.r.t) *m*^*th*^ annotation is

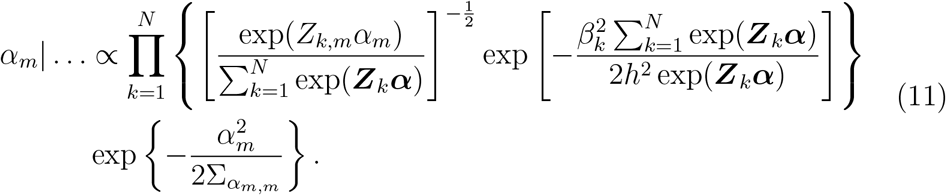

t-candidate distribution is used in MH sampler with location parameter obtained by maximizing ***α***_*m*_ over equation (11).

### 9.2 Simulations

#### 9.2.1 Model Performance

In order to evaluate the model performance, a matrix ***X*** of dimension 9,323 *×* 46,360 (n:N is approximately 1:5) is obtained from ABCD study by taking features (cortical vertices) from baseline, if missing, then from 2-year visit. The difference between this ***X*** and the ***X*** described in Data Section (5) lies on xxxxxx. 10 binary annotations are utilized in the simulation: Default CinguloOperc, DorsalAttn CinguloOperc, SMhand CinguloOperc, Visual CinguloOperc, Default Default, DorsalAttn Default, SMhand Default, Visual Default, Visual DorsalAttn and Visual SMhand. The selection of annotations is not important here; what important is that they are realistic feature-level annotations.

Next, columns of features in ***X*** are extracted based on the union of value 1’s across 10 selected annotations resulting in the subset of the features *Ñ* = 14, 019. Subsequently, 2,804 (*ñ*) rows from the full feature matrix has been randomly selected yielding the ratio of *ñ* : *Ñ* from submatrix 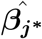 remaing 1:5. As for model parameters, true value of *h*^2^ is set to 0.3 representing FVE (fraction of the variance explained) value; ***α*** is set to 1.5 and -0.5 to represent large and small annotation effects, respectively. All 10 ***α*** values (mapping to 10 annotations) are set to 1.5 in one batch and -0.5 in the other. The main purpose of the simulations is to examine if the parameter estimates, especially ***α*** and *h*^2^, can converge to their true values with different annotations and different random selections of rows from ***X. β*** is drawn from distribution and 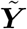 is generated from distribution (2) described in Section (3.1).

MCMC is a stochastic process with random nature, running multiple chains and average results across chains can reduce bias in estimates. For each of the two batches, Bayesian model inference is implemented one annotation per chain (e.g, *M* = 1, single-annotation model) and 10 chains per batch. The output of summary statistics from two batches are displayed in Table 3. For each batch, median values of estimates from the 10 MCMC chains are collected, the average and (minimum, maximum) of the 10 medians are reported. Although 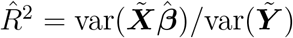 is not a parameter in the Bayesian model, it is the variance explained by the features directly from the regression and it is closely related to *h*^2^. As we can see, 95% posterior credible interval (CI) estimates of 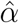 and *ĥ*^2^ cover the true values in both batches, *ĥ*^2^ and 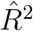 turn out to be very similar. 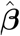 is a vector estimate, thus 95% posterior CI coverage rate of true value from each chain is calculated, then the average and (minimum, maximum) coverage rates from the 10 chains are reported for both batches.

**Table 3:** Simulation Summary Statistics.

| Estimates<sup>1</sup> | Batch 1 | Batch 2 |
| --- | --- | --- |
| | $\alpha = 1.5^*, h^2 = 0.3^*$ | $\alpha = -0.5^*, h^2 = 0.3^*$ |
| $\hat{\alpha}$ | 1.41 (1.15,1.65) | -0.60 (-1.34,-0.09) |
| $\hat{h}^2$ | 0.29 (0.26,0.33) | 0.30 (0.27,0.35) |
| $\hat{R}^2$ | 0.27 (0.22,0.32) | 0.29 (0.24,0.32) |
| ${}^2\hat{\beta}$, rate of covering true value | 97% (91%,99%) | 95% (92%,97%) |

<sup>1</sup>Median values of $\hat{\alpha}$ , $\hat{h}^2$ and $\hat{R}^2$ from the 10 MCMC chains per batch are collected, the average and (minimum, maximum) of the 10 medians are reported. $\hat{R}^2 = \text{var}(\tilde{\mathbf{X}}\hat{\beta})/\text{var}(\tilde{\mathbf{Y}})$ .
<sup>2</sup> $\hat{\beta}$ is a vector estimate, 95% posterior CI coverage rate of true value from each chain is calculated first, then the average and (minimum, maximum) coverage rates of the 10 chains per batch are reported.
\*True values set in the simulations.

**Table 4:**
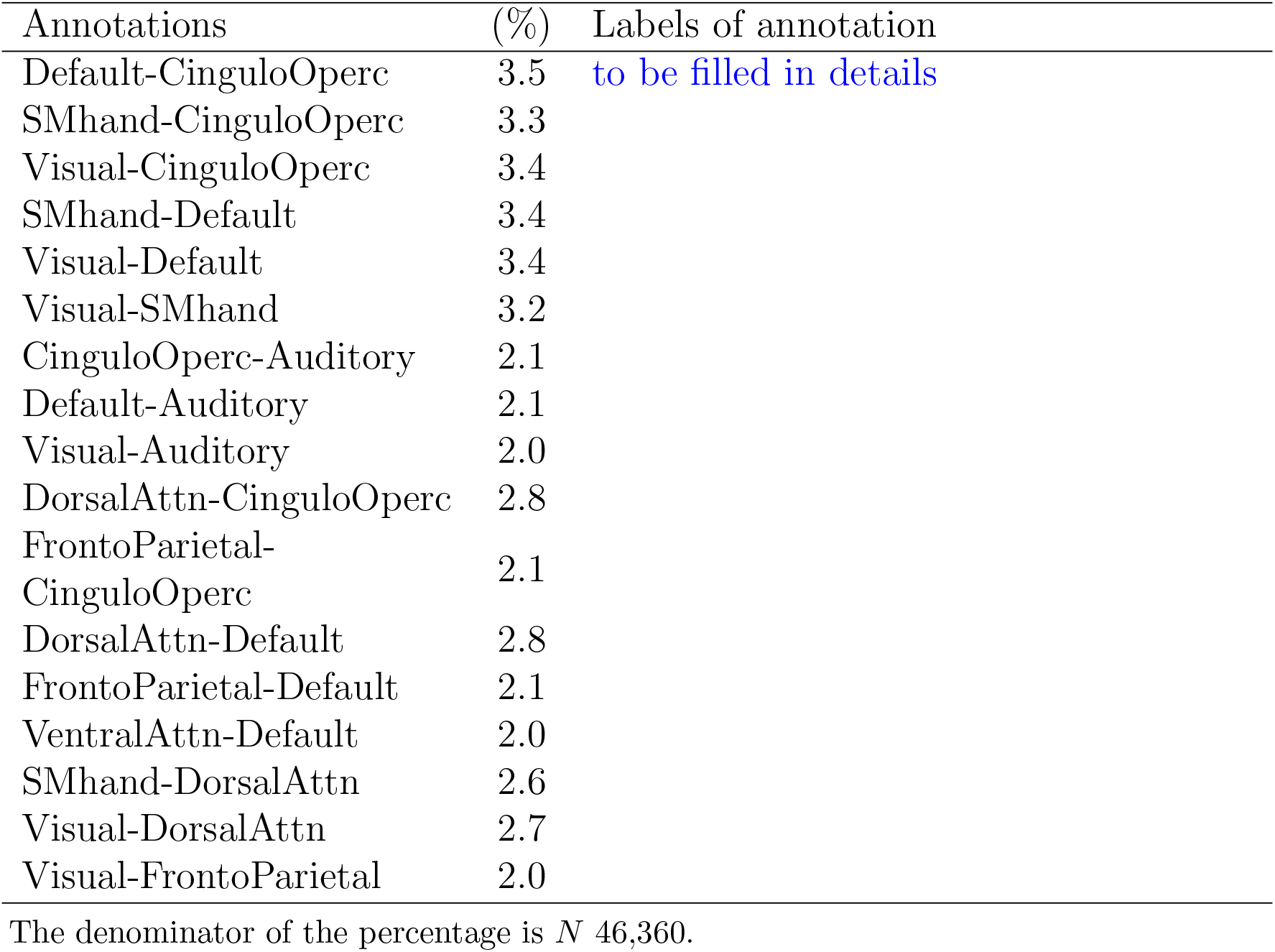
Percentage of 1’s from 17 between-network binary annotations.

**Table 5:**
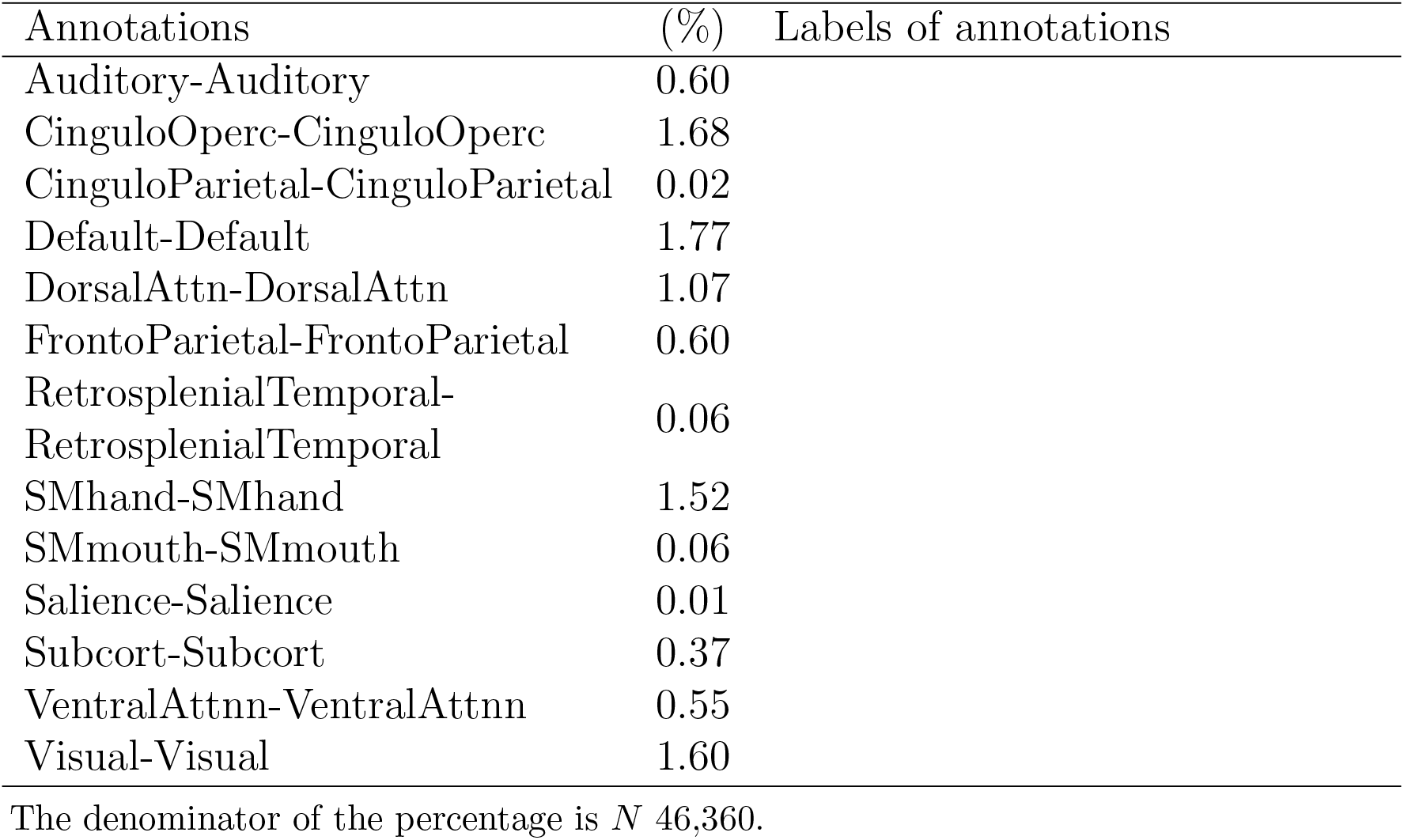
Percentage of 1’s from 13 within-network binary annotations.

#### 9.2.2 Validation of out-sample *h*^2^ estimate

To evaluate the out-sample performance of *h*^2^, the prior of ***β*** from (3) is simplified to MVN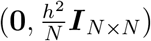 without annotations.

In first simulation, the true value of *h*^2^ is set to 0.32 and the true value of ***β*** is generated from the simplified prior distribution. The feature matrix ***X*** is randomly split into training set (***X***_*tr*_) and test set (***X***_*ts*_) 10 different times to create 10 training sets and 10 test sets. Each time training set sample size is 8,665 and test set is 320; ***X***_*tr*_ and ***X***_*ts*_ are then residulized by few subject-level covariates to eliminate any potential influence: age, female, MRI device serial number, genetic ancestry factors (African, East Asian and American),rsfmri_var_mean.motion and MRI manufacturers model name (some descriptions of these covariates?). This is the main reason that the total sample size in this simulation (8,665+320=8,985) is smaller than those from previous Simulation Section (9.2.1), because of the missing values in the covariates. Both ***X***_*tr*_ and ***X***_*ts*_ are standardized afterwards, respectively. Outcome variable, ***Y*** _*tr*_ and ***Y*** _*ts*_, are then generated from model (2) based on standardized ***X***_*tr*_ and ***X***_*ts*_, respectivly, as well as true values of *h*^2^ and **β**.

Bayesian model inference is applied to the 10 training sets to obtain parameter estimates 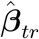 and 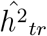, which are the median values of MCMC chains. In-sample estimate 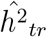 is ranged from 0.30 to 0.36 indicating reliable and stable estimates of *h*^2^. The squared correlation between 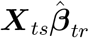 and ***Y*** _*ts*_ represents the out-sample *h*^2^ estimate from the test sets, which is ranged from 0.10 to 0.21 with median value of 0.15.

The procedure of the second simulation is very similar to the first one with few exceptions: (1) the dimension of ***X*** matrix is 5, 721 *×* 6, 1776, which is aligned with the ***X*** from Data Section (5); it has fewer subjects and more features, the n:N is approximately 1:11. (2) ***X*** is randomly split into training set (***X***_*tr*_) and test set (***X***_*ts*_) 6 different times to create 6 training sets and 6 test sets (due to the limitation of server capacity). Each time training set sample size is 5,401 and test set is 320. (3) subject-level covariates that ***X***_*tr*_ and ***X***_*ts*_ are regressed on: age, sex, MRI device serial number, MRI softwareversion, C1, C2, C3, C4, C5, C6, C7, C8, C9, C10. (4) The true value of *h*^2^ is set to 0.1. The results show that in-sample estimate 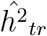 is ranged from 0.06 to 0.15. The out-sample estimate is ranged from 0.01 to 0.04 with median value of 0.03.

### 9.3 Computational Resource

The algorithm has been implemented in Python version 3.7.3 [9, 12] with multiprocessing invoked to run multiple MCMC chains simultaneously. The simulations are run on a Linux server with total 250 gigabytes (GB) of RAM and 20 CPUs that is dual Intel(R) Xeon(R) E5-2630 v4 2.20GHz. In simulations of model performance with 14,019 features, a chain of 30,000 iterations (10,000 burn-in, thinning of 10) takes about 16 hours; in simulations of *ĥ*^2^ out-sample validation with 46,360 features, a chain of 10,000 iterations (3,000 burn-in, thinning of 10) takes about 63 hours. The real data applications of ABCD are run on Linux server XXXX, with ***X*** dimension of 9, 323 *×* 46, 360, 10,000 iterations (3,000 burn-in, thinning of 10) takes about XX hours.

### 9.4 Binary Annotation (%)

